# Chronic deep brain stimulation of the human nucleus accumbens region disrupts the stability of inter-temporal preferences

**DOI:** 10.1101/2020.12.11.417337

**Authors:** Ben J. Wagner, Canan B. Schüller, Thomas Schüller, Juan C. Baldermann, Sina Kohl, Veerle Visser-Vandewalle, Daniel Huys, Milena Marx, Jens Kuhn, Jan Peters

## Abstract

When choosing between rewards that differ in temporal proximity (inter-temporal choice), human preferences are typically stable, constituting a clinically-relevant transdiagnostic trait. Here we show in patients undergoing deep brain stimulation (DBS) of the anterior limb of the internal capsule / nucleus accumbens region for treatment-resistant obsessivecompulsive disorder (OCD), that long-term chronic (but not phasic) DBS disrupts inter-temporal preferences. Hierarchical Bayesian modeling accounting for temporal discounting behavior across multiple time points allowed us to assess both short-term and long-term reliability of inter-temporal choice. In controls, temporal discounting was highly reliable, both long-term (6 months) and short-term (1 week). In contrast, in patients undergoing DBS, short-term reliability was high, but long-term reliability (6 months) was severely disrupted. Control analyses confirmed that this effect was not due to range restriction, the presence of OCD symptoms or group differences in choice stochasticity. Model-agnostic between- and within-subject analyses confirmed this effect. These findings provide initial evidence for long-term modulation of cognitive function via DBS and highlight a potential contribution of the human nucleus accumbens region to inter-temporal preference stability over time.

**Significance Statement:** Choosing between rewards that differ in temporal proximity is in part a stable trait with relevance for many mental disorders, and depends on prefrontal regions and regions of the dopamine system. Here we show that chronic deep brain stimulation (DBS) of the human anterior limb of the internal capsule / nucleus accumbens region for treatment-resistant obsessive compulsive disorder disrupts the stability of inter-temporal preferences. These findings show that chronic stimulation of one of the brain’s central motivational hubs can disrupt preferences thought to depend on this circuit.

## Introduction

Humans continuously maneuver their world weighting the future against the present. The degree of temporal discounting of future rewards (as assessed via inter-temporal choice tasks) is at least in part a stable trait (Kirby, 2009) with relevance for many mental disorders (Amlung et al., 2019). For example, steep discounting of value over time is a hallmark of substance use disorders (Amlung et al., 2017; Bickel et al., 2014), gambling disorder (Dixon et al., 2003; Wiehler & Peters, 2015) and a range of other disorders (Amlung et al., 2019; Lempert et al., 2019).

Multiple neural systems contribute to human self-control, including prefrontal cortex (PFC) regions involved in cognitive control, and regions of the mesolimbic and mesocortical dopamine system (e.g. ventral striatum and ventro-medial PFC) involved in reward valuation (Bartra et al., 2013; McClure & Bickel, 2014; Peters & Büchel, 2011). Earlier studies focused on characterizing potentially dissociable striatal and prefrontal value signals during inter-temporal choice (Kable & Glimcher, 2007; McClure et al., 2004). This debate has ultimately led to a revised view of self-control, according to which lateral PFC exerts top-down control over ventromedial PFC in support of self-controlled choices (Figner et al., 2010; Hare et al., 2009; Peters & D’Esposito, 2016), where both striatal reward responses (Hariri et al., 2006) and cortico-striatal connectivity (van den Bos et al., 2014) are associated with temporal discounting in cross-sectional analyses. Despite these findings and direct evidence for a contribution of the ventral striatum to intertemporal choice in rodents (Pisansky et al., 2019), direct evidence in humans is lacking.

At the same time, deep brain stimulation (DBS) to different subcortical target sites has gained increasing attention in psychiatry in the context of different mental disorders (Krauss et al., 2021; Lee et al., 2019). This includes e.g. depression (Dandekar et al., 2018), obsessive compulsive disorder (OCD) (Kuhn & Baldermann, 2020; Wu et al., 2020), substance use disorders (Mahoney et al., 2020; Peisker et al., 2018) and schizophrenia (Nucifora et al., 2019). While clinical outcomes are in many cases promising, one oftentimes neglected issue concerns potentially more subtle cognitive effects that might arise as a result of chronic stimulation.

Here we address this issue by capitalizing on the rare opportunity to longitudinally follow patients undergoing therapeutic deep brain stimulation (DBS) of the anterior limb of the internal capsule / nucleus accumbens (ALIC/NAcc) region for treatment-resistant OCD. OCD is assumed to be associated with a dysregulation in fronto-striatal circuits (Robbins et al., 2019), which might be normalized via ALIC/NAcc DBS (Figee et al., 2013; Smith et al., 2020; Smolders et al., 2013; Wu et al., 2020). In the context of a DBS treatment-efficacy study (Huys et al., 2019) we comprehensively examined short-term (acute) and long-term (chronic) effects of DBS on intertemporal preference. Analyses focused on three aspects. First, we analyzed group differences in temporal discounting between patients with OCD and controls, which previously yielded heterogeneous results. Second, we directly tested for effects of acute DBS on inter-temporal choice by comparing ON vs. OFF stimulation periods in patients. Finally, effects of long-term (i.e. chronic) DBS on inter-temporal choice were examined by comparing long-term reliability estimates of temporal discounting between controls and patients undergoing DBS. These analyses leveraged hierarchical Bayesian parameter estimation using generative models to directly estimate posterior distributions of test-retest reliabilities, separately for each group, and were corroborated by model-agnostic control analyses.

## Materials and Methods

### Participants

All participants provided written informed consent prior to participation, and the study procedure was approved by the Ethics Committee of the Medical Faculty of the University of Cologne. Table 1 provides an overview of demographics data for patients and controls.

**Table 1.** Demographic data per group. Scores are Mean (SD).

| Long-term reliability | Controls (n = 28) | Patients (n = 9) | Group Comparison |
| --- | --- | --- | --- |
| Age (years) | 40.2 (9.0) | 41.4 (11.6) | $t_{(11.223)} = -0.30, p = 0.77$ |
| Sex (F/M) | 14/14 | 4/5 | $X^2_{(1)} < 0.001, p = 1$ |
| Education (years) | 11.9(1.4) | 10.7(1.4) | $t_{(13.325)} = 2.34, p = 0.04$ |
| Short-term reliability | Controls (n = 27) | Patients (n = 15) |  |
| Age (years) | 40.1 (9.1) | 47.4 (11.3) | $t_{(21.944)} = -2.08, p = 0.05$ |
| Sex (F/M) | 13/14 | 8/7 | $X^2_{(1)} = 0, p = 1$ |
| Education (years) | 11.9 (1.4) | 10.8 (1.5) | $t_{(24.4)} = 2.26, p = 0.03$ |
| Long-term (β-matched) | Controls (n = 14) | Patients (n = 9) | |
| Age (yrs) | 41.6 (9.9) | 41.4 (11.6) | $t_{(14.305)} = .044, p = 0.97$ |
| Sex (F/M) | 9/8 | 5/4 | $X^2_{(1)} < 0.001, p = 1$ |
| Education (yrs) | 11.35 | 10.7(1.4) | $T_{(27.577)} = 0.96, p = 0.34$ |

### Overall procedure

Patients with OCD and matched controls (for inclusion criteria and demographics see below) completed three separate testing sessions, where they completed a temporal discounting task (see below). One initial testing session (T1) corresponded to the pre-DBS session in patients, and the first testing session in controls. This was followed by two follow-up sessions at T2, T2.1 and T2.2. The interval between T1 and T2 sessions in days was matched between groups (M [range] patients: 203 [155-260], controls: 206 [164-247]). The T2.1 and T2.2 sessions were spaced within a week. Here, patients performed DBS on vs. off sessions in counterbalanced order with at least 24h wash-out period, whereas controls performed simply two addition testing sessions. A subset of patients (n=7) completed all three testing sessions. Two additional patients completed T1 but only one of the two T2 sessions, yielding a total n=9 for pre vs. post DBS analyses in patients. Eight additional patients were only tested at T2, yielding a total of n=15 for on vs. off DBS analyses.

In total, n=28 controls participated (see below). One control participant missed the final testing session, yielding n=27 for the corresponding analyses. The Covid-19 pandemic broke out while this study was being conducted. Therefore, ten of the controls completed testing at T2 online, as opposed to in the lab (see below for control analyses).

### Patients with OCD

Patients with OCD eligible for DBS had to meet the DSM-IV criteria for OCD, a Yale-Brown Obsessive Compulsive Scale (Y-BOCS) over 25, at least one cognitive-behavioral therapy (minimum of 45 sessions), at least two unsuccessful treatments with a serotonin reuptake inhibitor and one unsuccessful augmentation with either lithium, neuroleptics or buspirone. Patients were excluded due to drug, medication or alcohol abuse, suicidal ideation, mental retardation, pregnancy or breastfeeding and schizophrenia. Disease duration was on average 27.59 ± 13.02 years ranging from 6 to 48 years. The mean age at onset for OCD was 16.2 ± 9.25 years. Individual patient clinical data are provided in Table 2.

**Table 2.**
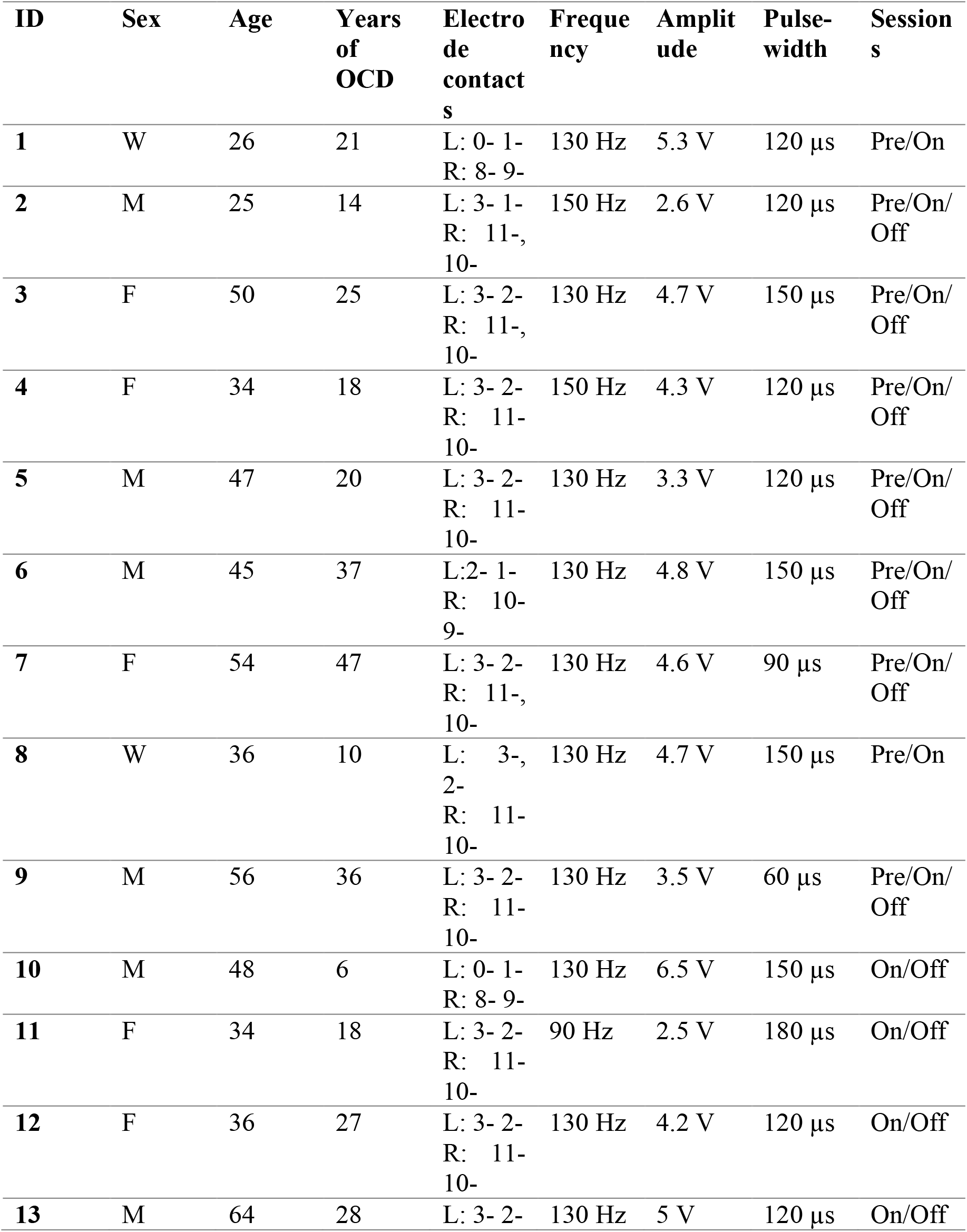
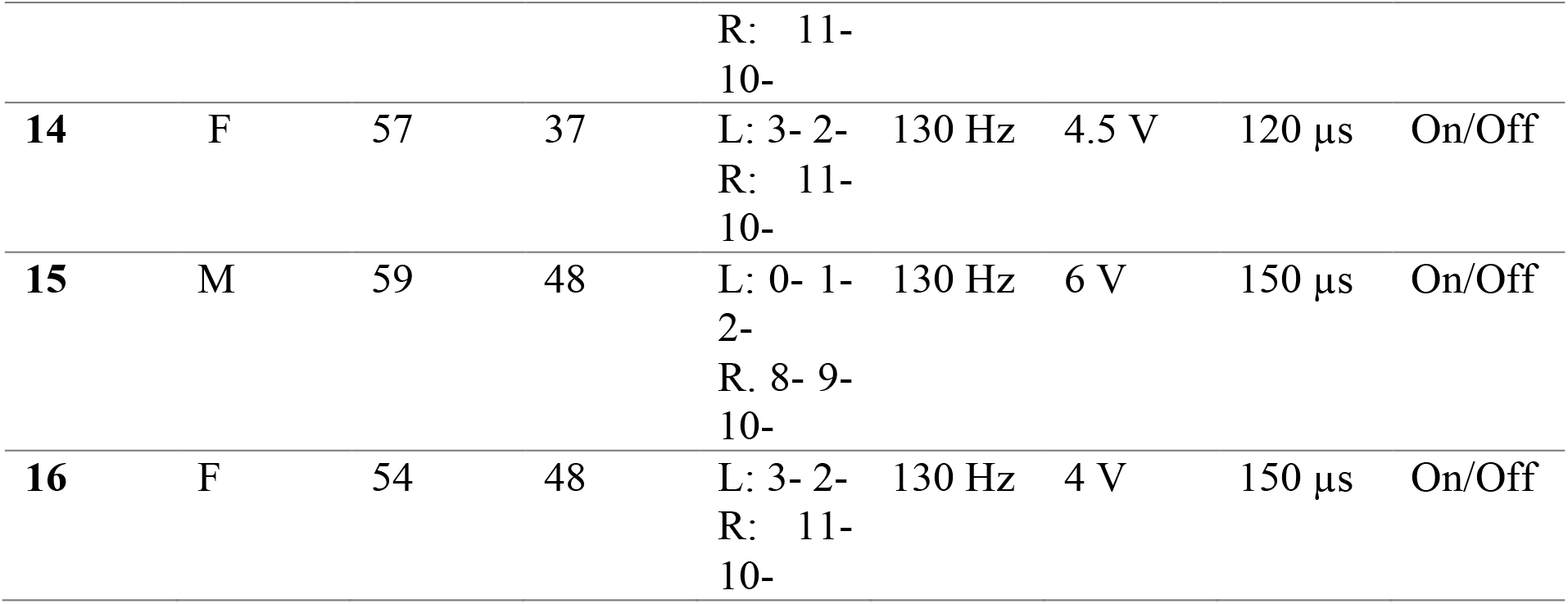
Overview of sex, age, disease duration before surgery and stimulation parameters (monopolar, all bilateral, except for patient 13) of DBS patients with OCD. DBS, deep brain stimulation; F, Female; Hz, Hertz; L, Left; M, Male; µ;s, microsec; OCD, obsessive-compulsive disorder; R, Right.

### Controls

In total, n=28 controls participated in the study (see Table 1 for demographic data). Exclusion criteria were drug, medication or alcohol abuse or the diagnosis of a psychiatric disorder. Controls were screened for OCD-symptoms via the OCI-R questionnaire (Foa et al., 2002). Here 9/28 subjects scored above the threshold (>21) for possible OCD.

### Temporal discounting task

Prior to the first testing session, participants completed a short adaptive pretest to estimate the individual discount-rate (*k*). This discount rate was used to construct a set of 140 participant-specific trials using MATLAB (version 8.4.0. Natick, Massachusetts: The MathWorks Inc). The primary task then consisted of choices between an immediate smaller-sooner reward of 20€ and participant specific larger-but-later (LL) rewards delivered after some delay (1, 2, 7, 14, 30, 90 or 180 days). In seventy trials, LL amounts were uniformly spaced between 20.5€ and 80€. In the remaining seventy trials, LL amounts were uniformly spaced around each estimated indifference point per delay (based on the pre-test discount rate). In case indifference points were > 80€, only uniformly-spaced LL amounts were used. Trials were presented in a pseudorandomized order. Following each task completion, one trial was randomly selected and paid out immediately in cash (in case of a smaller-sooner choice) or via a timed bank transfer (larger-but-later choice).

### Deep brain stimulation

DBS was applied to the ALIC/NAcc region. Details on electrode placement (including reconstruction of electrode positions), surgical procedure and adjustment of stimulation parameters are available elsewhere (Huys et al., 2019). Final stimulation amplitudes ranged from 2.6 to 4.8 volt and pulse-width was set between 60 and 150 µs (see Table 2). The frequency of DBS was 130 Hz except for two patients who received 150 Hz stimulation and one patient who received 90 Hz stimulation.

### Computational Modeling

To assess inter-temporal choice, we applied a standard single-parameter hyperbolic discounting model:

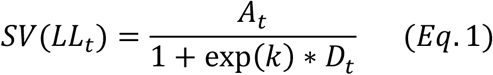

Here, *A* is the numerical reward amount of the *LL* option on trial *t*. The discount-rate (*k)*, here modeled in log-space, reflects the steepness of the hyperbolic discounting curve, with greater values corresponding to steeper discounting. Delay *D* of the LL option is expressed in days. *SV* then corresponds to the subjective (discounted) value of the delayed (LL) option. We then used softmax action selection (Sutton & Barto, 1998) (Eq. 2) to model the probability of selecting the *LL* option on trial *t*. Here, *β* is an inverse temperature parameter, modeling choice stochasticity. For *β* = 0, choices are random, and as *β* increases, choices become more dependent on option values:

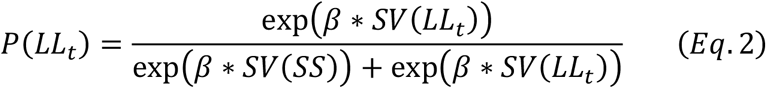

In a first step, models were fit to all trials from all participants, separately per group and time point, using a hierarchical Bayesian modeling approach. We included separate group-level Gaussian distributions for log(*k*) and *β* for patients and controls, and/or T1 and T2 or within T2 (T_2.1_, T_2.2_) time points (see below for information on prior distributions).

Secondly, we implemented a fully generative approach by drawing discount-rate parameters from multivariate Gaussian distributions, to obtain posterior distributions for the reliability (covariance). That is, for the case of two time points T1 and T2, log(*k*) values for a given participant *i* are drawn from a bivariate Gaussian:

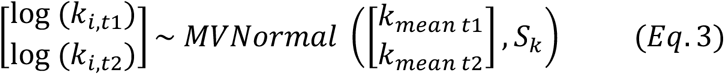

with variance-covariance matrix S_k_:

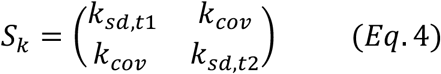

The diagonal contains the variances and the off-diagonal elements contain the covariances. The test-retest correlation coefficient *ρ* is directly estimated using:

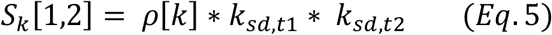

With

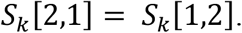

Separate models were used to estimate long-term reliability (T1/pre DBS and T2/post DBS in controls and patients respectively) and short-term reliability (i.e. T2.1/DBS on and T2.2/DBS off), separately in patients and controls.

### Hierarchical Bayesian parameter estimation

Parameter estimation was performed using Markov Chain Monte Carlo as implemented in the JAGS software package (Plummer, 2003) (Version 4.3) in combination with R (Version 3.4) and the R2Jags package. For group-level means, we used uniform priors defined over numerically plausible parameter ranges ([-20, 3] for log(*k*); [0, 10] for *β* and [-1,1] for *ρ*). We initially ran two chains with a varying burn-in period of at least 20.000 samples and thinning of two. Chain convergence was then assessed via the Gelman-Rubinstein convergence diagnostic 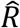 and sampling was continued until 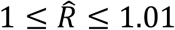 for all group-level and individual-subject parameters. 10k additional samples were then retained for further analysis. We report posterior group distributions for all parameters of interest as well as their 85% and 95% highest density intervals (HDIs). For group comparisons (patients vs. controls) and time effects (T1 vs. T2) we report Bayes Factors for directional effects for the hyperparameter difference distributions, estimated via kernel density estimation using R (Version 4.01) and RStudio (Version 1.3).

### Analysis of group differences

We report posterior difference distributions and the corresponding 85 % and 95 % highest density intervals to characterize differences between patients and controls, changes from T1 to T2 or within T2 (T_2.1_, T_2.2_; on/off DBS).. We then estimate Bayes Factors for directional effects. These were computed as the ratio of the integral of the posterior difference distribution from 0 to +∞ vs. the integral from 0 to -∞. Using common criteria (Beard et al., 2016), we considered Bayes Factors between 1 and 3 as anecdotal evidence, Bayes Factors above 3 as moderate evidence and Bayes Factors above 10 as strong evidence. Bayes Factors above 30 and 100 were considered as very strong evidence.

### Model-based choice inconsistency

For both model-based and model-agnostic within-participant changes, we leveraged the fact that participants completed the exact same 140 choices during each testing session. To examine model-based changes in preferences, we extracted individual-participant median discount-rates log(*k*) and decision noise parameters (softmax *β*) from our hierarchical Bayesian model estimated on T1 data. We then used these parameters to compute choice probabilities for each T1 choice. To examine model-based preference changes from T1 to T2, we then subtracted the T1 choice probability from the corresponding observed choices at T2 (0 for smaller-sooner and 1 for larger-later choices). We then computed a choice inconsistency score as the mean of the absolute differences between T1 choice probabilities and T2 choices.

Because controls showed lower decision noise (*β*) compared to patients, which was also reflected in an overall somewhat lower model fit in patients (Table 3), for additional control analyses, we also matched groups on *β*.

**Table 3.**
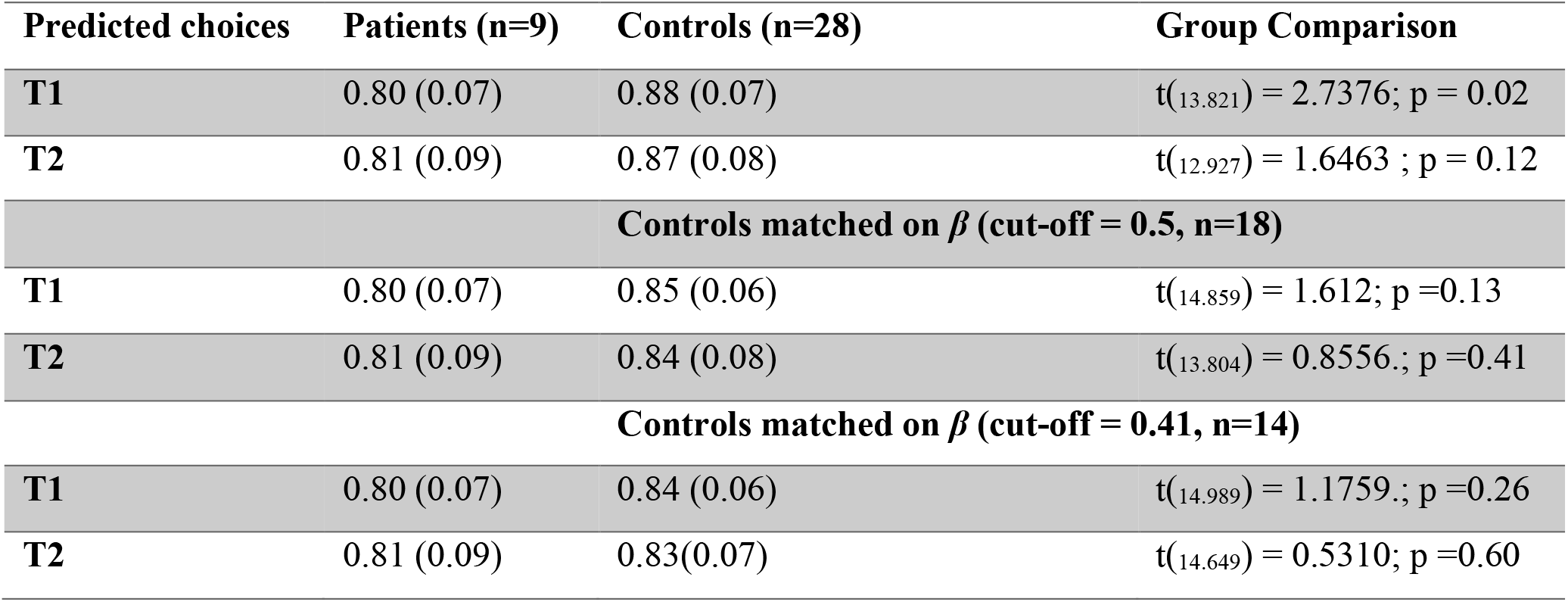
Mean (SD) proportion of correctly predicted choices of the full hierarchical Bayesian model (hyperbolic discounting + softmax + correlation in ρ log[*k*] and ρ softmax *β*) that jointly estimates discount-rate, softmax beta and the intraindividual correlation of both parameters. At T1 the model performed better in controls than in patients. This difference was not significant at T2. This difference also disappeared when groups were matched on decision noise (softmax *β*).

### Model agnostic analysis of indifference points

Model-based analyses rely on specific mathematical assumptions regarding the shape of the discounting function. Furthermore, they can be affected by potential group differences in model fit. Therefore, we additionally examined a model-agnostic measure of within-participant changes in preferences. To this end, we fit sigmoid functions (see Eq. 3) to the choice data of each delay *D* per participant and time point *T*:

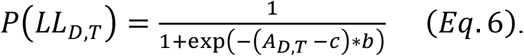

We modeled the probability to choose the delayed reward for delay D at time point T for each participant as a sigmoid function of the absolute LL reward amount *A*. Here, *c* is the inflection point of the sigmoid (corresponding to the indifference point, i.e. the point of subjective equivalence between the delayed reward and the immediate reward at the respective delay D), and *b* is the slope.

For delays with only larger-later choices, the indifference point was set to the midpoint between the immediate reward (20€) and the smallest available LL reward. For delays with only smaller-sooner choices, the indifference point was conservatively set to max(*LL*). These rules were also applied in cases where there was only a single noisy LL or SS choice for a given delay. Using this procedure, we computed 196 indifference points in controls and 63 indifference points in patients. Six indifference points in controls and two in patients with OCD could not be estimated. We then computed the mean absolute deviation in indifference points between T1 and T2 as a model-agnostic measure of preference consistency.

### Permutation-based group comparisons

Model-based and model-agnostic consistency measures (see previous sections) were compared between groups using permutation tests. To this end, we compared the observed group difference in preference consistency to a null-distribution of preference consistency that was obtained by randomly shuffling group labels 10k times, and computing the group difference for these shuffled data. Significance was assessed using a two-tailed *p*-value of 0.05.

### Code and data availability

Model code is available on the Open Science Framework (https://osf.io/53mkd/). Raw choice data are available from Zenodo.org (https://doi.org/10.5281/zenodo.7559218) for researchers meeting the criteria for access to confidential data.

## Results

Single-subject choice data, separately for T1 and T2 sessions, are shown in Figure 1 for individual OCD patients, and in Figure 2 for individual control participants.

**Figure 1.**
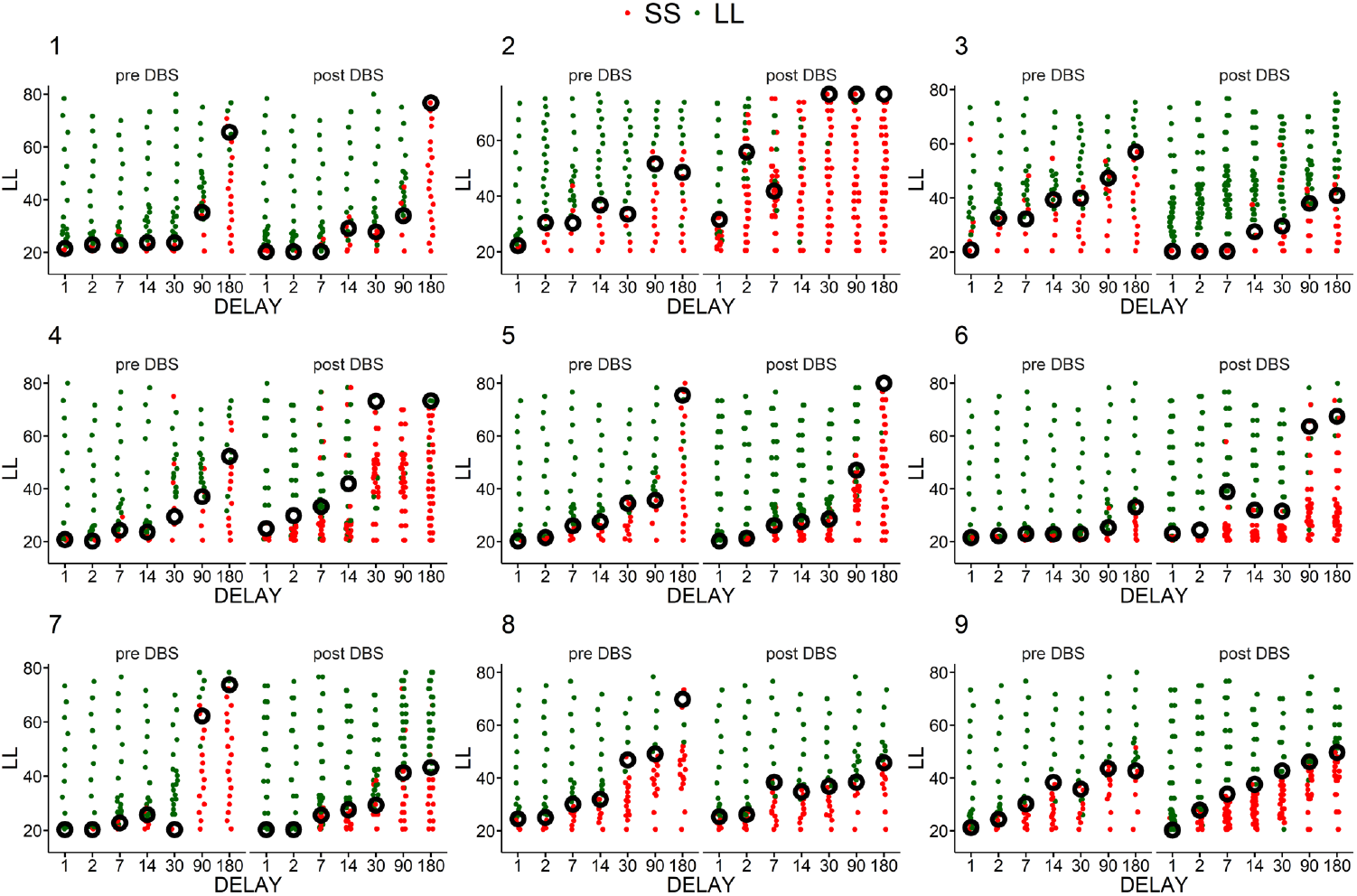
Single-subject choice data for all n=9 patients with pre- (T1) and post-DBS data (T2). Green and red points represent LL and SS choices, respectively, across LL amounts (y-axis) and delays (x-axis). Black circles show estimated indifference-points.

**Figure 2.**
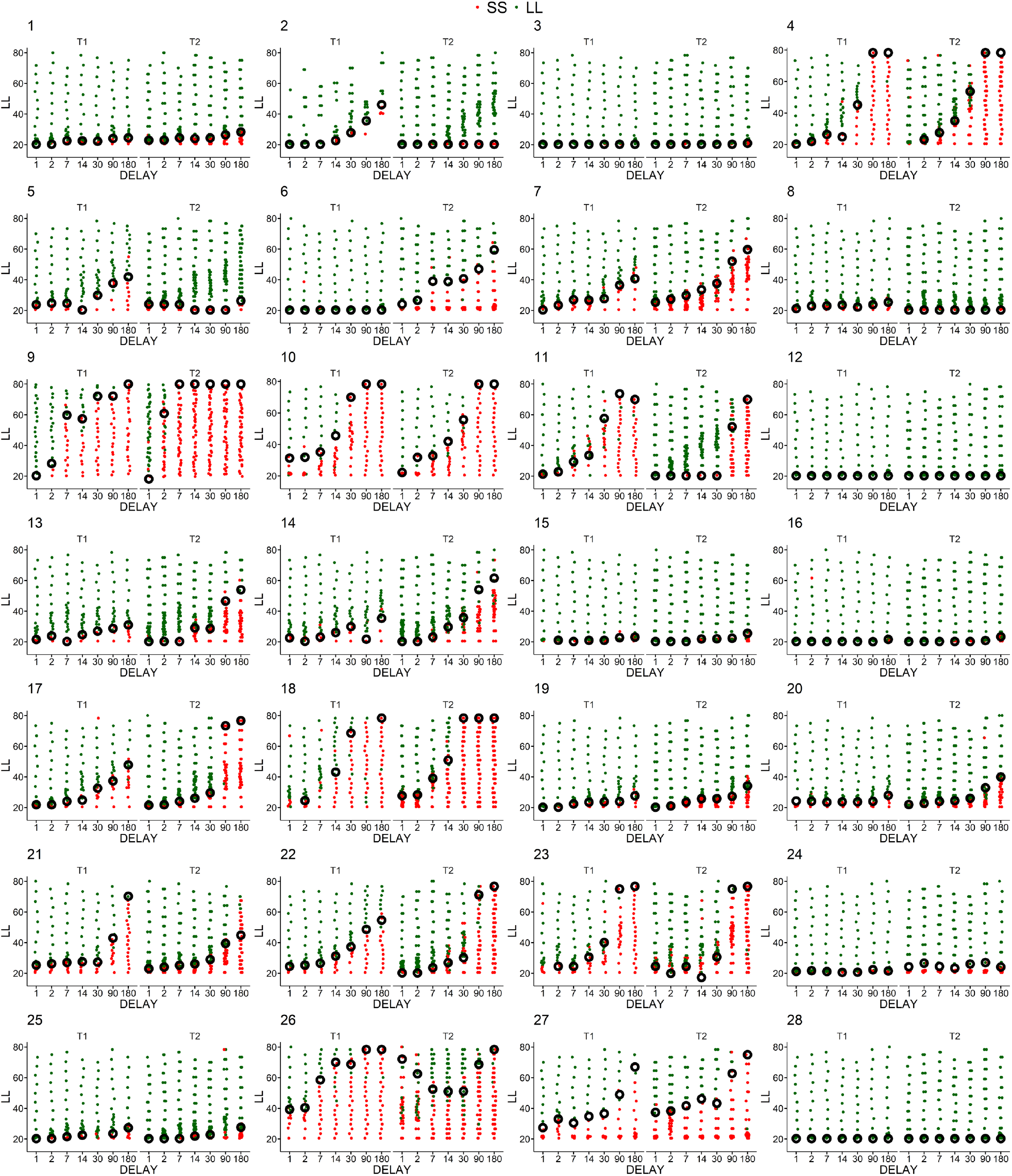
Single-subject choice data for all n=28 controls. Green and red points represent LL and SS choices, respectively, across LL amounts (y-axis) and delays (x-axis). Black circles show estimated indifference-points.

In a first step, data were then modeled for each time point separately using hierarchical Bayesian parameter estimation using a hyperbolic discounting model with softmax action selection. This model accounted for ≥ 80% of choices in both patients and controls across all testing sessions (see Table 3).

In line with previous work (Sohn et al., 2014), patients with OCD exhibited increased discounting (a higher discount rate log[*k*]), both pre DBS at T1 (Figure 3A; the directional Bayes Factor (dBF) for a group difference <0 was 35.57; 95% HDI of the difference distribution: -2.28 – 0.04) and post DBS at T2 (Figure 3B; dBF [< 0] = 15.85; 95% HDI = - 2.95 – 0.34).

**Figure 3.**
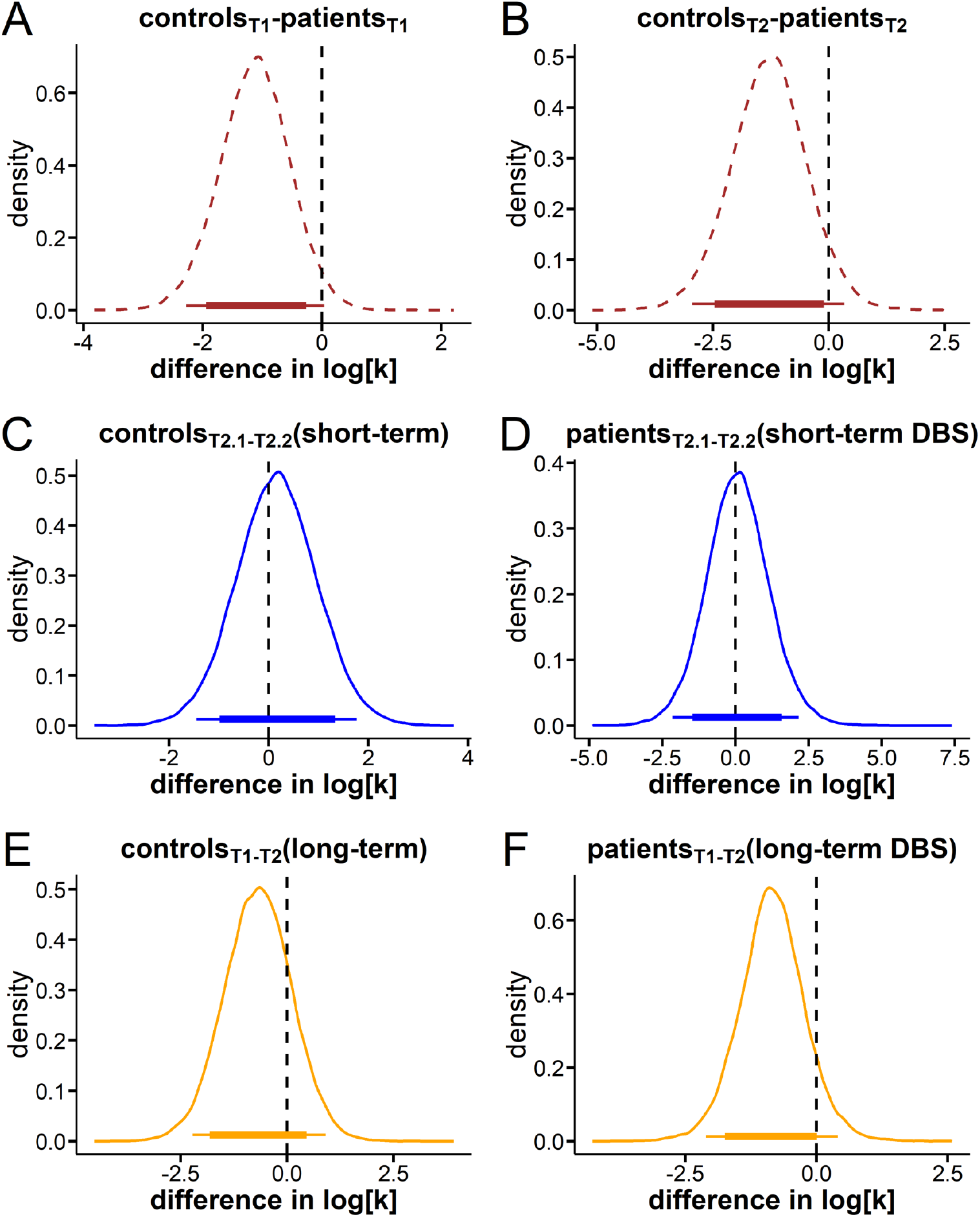
Group-level effects in inter-temporal choice. **A**: At T1 (pre DBS), partients (n = 9) discounted delayed rewards steeper compared to controls (n = 28) (directional Bayes Factor (dBF): controls < patients = 35.75; HDI = -2.28 – 0.04). **B**: Comparison of pooled T2 sessions in controls (n = 28, T2.1 and T2.2) and pooled DBS on and DBS off sessions at T2 in patients (n = 9). Higher log[k] values in controls were around 15.85 times more likely than lower values (95% HDI of the difference distribution: -2.95 – 0.34). **C**: There was no evidence for a systematic change in log[*k*] across T2 sessions in controls (dBF = 0.73). **D**: Likewise there was no systematic effect of acute DBS at T2 in patients (on. vs. off, n = 15, dBF = 0.90). **E** and **F**: If anything, both controls (E, n = 28) and patients (F, n = 9) tended to discount rewards steeper after six months (controls: T1 < T2; dBF = 4.15; 95% HDI = -2.22 – 0.90; patients: pre DBS < post DBS; dBF = 10.87; 95% HDI = -2.10 – 0.41). All panels depict posterior distributions smoothed via kernel density estimation in R. Thin (thick) horizontal lines denote the 95% (85%) highest density intervals.

Controls and patients showed no systematic short-term change in discounting (log[*k*]) between the two testing sessions at T2 (see Figure 3 C and D). Since this comparison corresponds to comparing on vs. off DBS sessions in patients, we found no systematic effect of phasic DBS on temporal discounting. With respect to the long-term changes in temporal discounting, if anything, rewards where discounted somewhat steeper after six months in controls (Figure 3E; dBF [<0] = 4.15; 95% HDI = -2.22 – 0.90) or following six months of continuous DBS in patients (Figure 3F; dBF [< 0] = 10.87; 95% HDI = -2.10 – 0.41), a pattern observed previously in healthy control participants (Kirby, 2009).

We applied both inter- and intra-individual analytical approaches to study DBS effects on the stability of inter-temporal preferences,. First, we used a generative modeling approach, modeling log[*k*] values across multiple time points using multivariate gaussian distributions (Haines et al., 2020) (see methods section). This allowed us to directly estimate a posterior distribution for the test-retest correlation ρ of log[*k*], separately for patients and controls, and for short-term (T2.1 vs. T2.2) and long-term (T1 vs. T2) reliability. In controls, the test-retest reliability of log[*k*] was high, both across T2 sessions (Figure 4A: one-week short-term reliability) and between T1 and pooled T2 data (Figure 4B, six-month long-term reliability). In both cases, the mean posterior distribution of the reliability coefficient ρ was ≥ .8, indicating high reliability. In patients, short-term reliability was likewise high (Figure 4D). In stark contrast, long-term reliability in patients was completely disrupted (Figure 4E). For short-term reliability in patients, the mean posterior distribution of the reliability coefficient ρ was ≥ .95, indicating excellent reliability. In contrast, for long-term reliability in patients, ρ was ≈ -.30, and the 95% HDI overlapped with zero (see Figure 4F).

**Figure 4.**
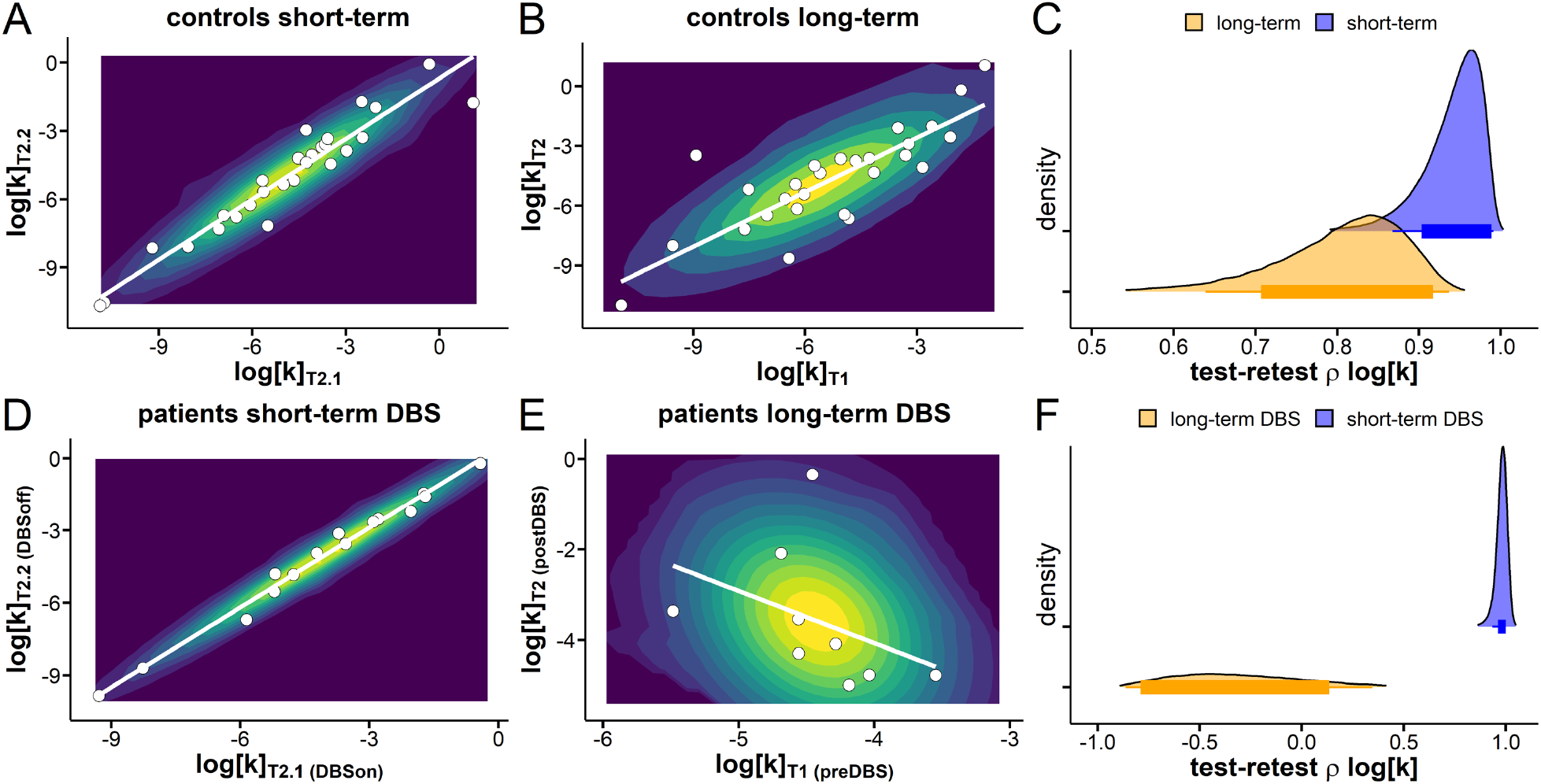
Reliability of inter-temporal choice (discount rate log[*k*]). **A**, and **B** show contour plots of the fitted multivariate gaussian and individual median discount-rate parameters log[*k*] for T1 and T2 sessions in controls (A,B) and patients (D, E). The x- and y axis with regard to the short-term data refer to the comparison within T2. **A**: control session T2.1 (x-axis) vs. T2.2 (y-axis). With regard to the long-term data (B, E) the axes refer to T1 (x-axis) vs. pooled T2.1 and T2.2 data (y-axis). **D**: T2.1 (DBS on) vs. T2.2 (DBS off) log[*k*] values in patients at T2. **E**, In the long-term condition data the x-axis shows discount-rates in the T1 (pre-DBS) session and the y-axis refers to the pooled T2 sessions (post DBS). **C**,**F**: ρ log[*k*] hyperparameter capturing the correlation of short- and long-term reliability of intertemporal preference in controls (**C**) and patients (**F**). All parameters were estimated from our hierarchical Bayesian model.

We then conducted a range of control analyses. First, to ensure that results were not affected by the different ranges of log[*k*] values in the two groups, we drew a subset of n=9 participants from the control group, matched on the exact range of log[*k*] values in patients. This subset of controls still exhibited significant long-term reliability (see Figure 5A, mean ρ = .55), and the 95% HDI of the posterior distribution of ρ was > 0. Second, to ensure that results are not affected by potential group differences in decision noise (*β*-parameter), we likewise drew a subset of n=14 controls matched to the patients on *β*. This subset of controls still exhibited significant long-term reliability (see Figure 5A, mean ρ = .77), and the 95% HDI of the posterior distribution of ρ was again > 0 (Figure 5A). Third, the loss of long-term reliability was also not generally due to the presence of OCD symptoms, as the n=14 controls with the highest OCI-R scores (in a range similar to some OCD patients e.g. Pinto et al., 2014) still exhibited significant long-term reliability (mean ρ = .94) with the 95% HDI again not overlapping with 0.

**Figure 5.**
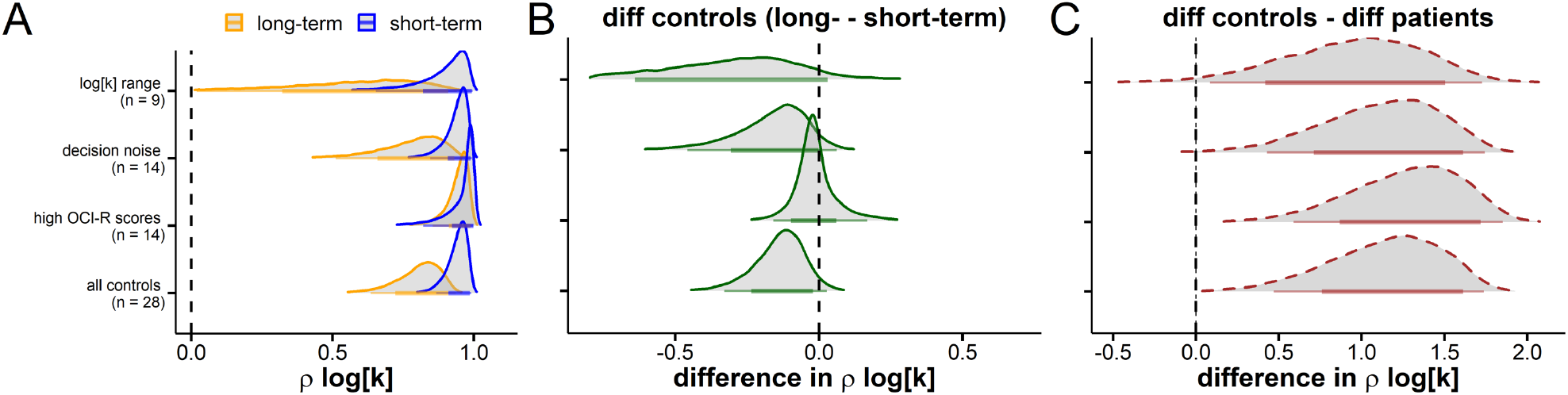
Control analyses. **A**, Posterior distributions of ρ log[*k*] (i.e. the test-retest reliability of the discount rate log[*k*]) for both short-term and long-term reliabilities and various subgroups of the control group. First row: controls matched on the range of log[*k*] values of the patient group, to account for effects of log[*k*] variance on reliability. Second row: controls matched on decision noise of the patient group to account for effects of decision-noise on reliability. Third row: controls with high OCI-R scores (mean[range] = 23.57 [14-40) to account for effects of OCD symptoms on reliability. Fourth row: all control participants. **B**, Difference between long-term and short-term reliability for all control subgroups. **C**, Group difference (controls vs. patients) in short-term vs. long-term reliability of all control subsamples. Here, values > 0 reflect lower long-term reliability in patients vs. controls.

To directly quantify group differences in reliabilities between controls and patients, we next computed differences in long- vs. short-term reliability posterior distributions for each group, and then compared these differences distributions across groups. Note that if group differences in long-term vs. short-term reliability are similar across groups, this difference distribution should not be different from zero. In contrast, if the disrupted long-term stability in patients is greater than what would be expected, given the control data, this difference distribution should be reliably greater than zero. In all control groups, long-term reliability was numerically lower than short-term reliability (Figure 5B), but the group difference in this difference was reliably > 0 in all cases (Figure 5C), reflecting a disrupted long-term reliability in the patient group.

Figure 6 summarizes these results for the full control sample (n=28). This shows a much more substantial difference in long vs. short-term reliability in patients (Figure 6A), resulting in a reliable group difference in reliabilities, with the 95% HDI of the difference distribution not overlapping with 0 (Figure 6B).

**Figure 6.**
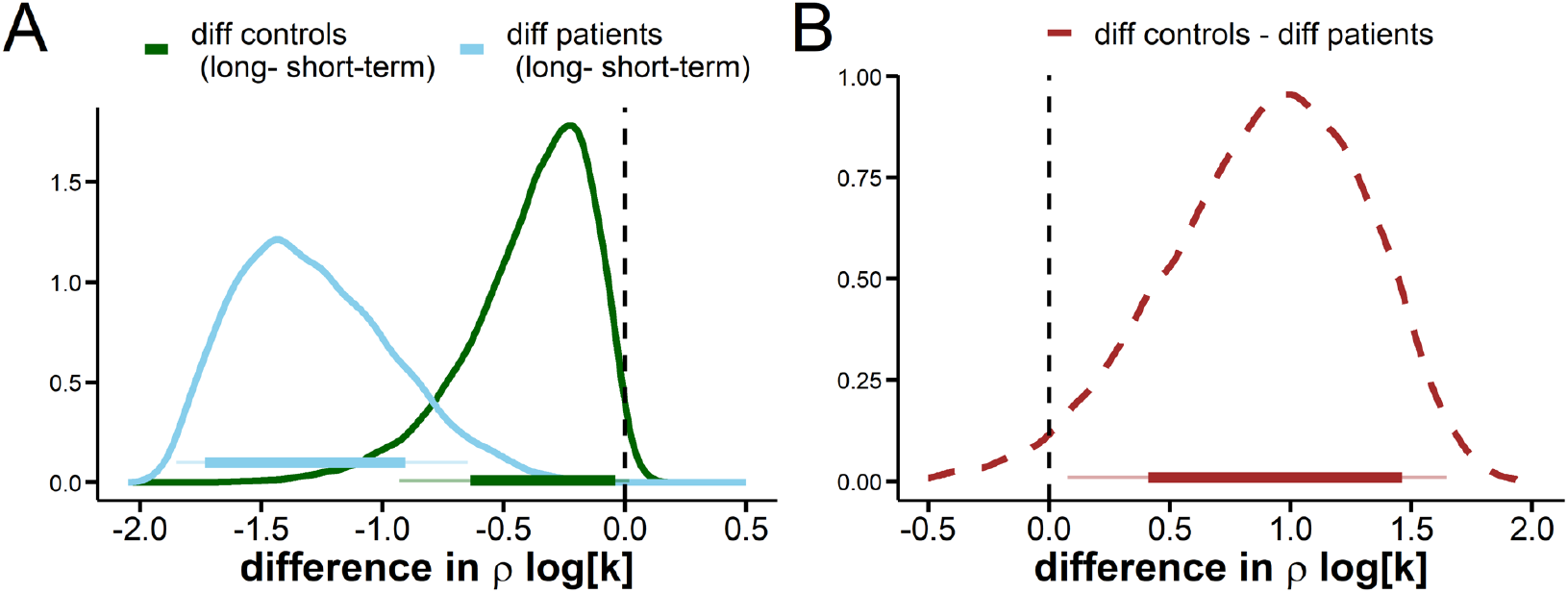
**A**, Difference in long- vs. short-term reliability (i.e. test-retest correlation coefficient *ρ* log[*k*]) in controls (green) and patients (blue). In controls, the 95% HDI overlapped with zero (95% HDI -.93 - .02), there was no overlap in patients. **B**, The group difference of the distributions shown in **A**. The 95% HDI did not overlap with zero (0.08 - 1.65; dBF = 36.78), indicating a reliable group difference.

In a last step, we tested whether a disruption of preference stability (as reflected in the disruption of long-term reliability) would also manifest at the level of individual decisions. Using both model-based and model-agnostic techniques (via indifference points and standard statistical analyses, see methods section) preference stability at the individual-participant level was quantified. As a model-based measure of stability, the mean deviation of T2 choice probabilities from predictions of a model fitted to T1 data was computed (Figure 7A). This inconsistency measure was greater in patients vs. controls (permutation test, Figure 7B). As a model-agnostic measure of stability, the mean absolute deviation of T1 and T2 indifference points was computed (Figure 7C). This inconsistency measure was again greater in patients vs. controls (permutation test, Figure 7D).

**Figure 7.**
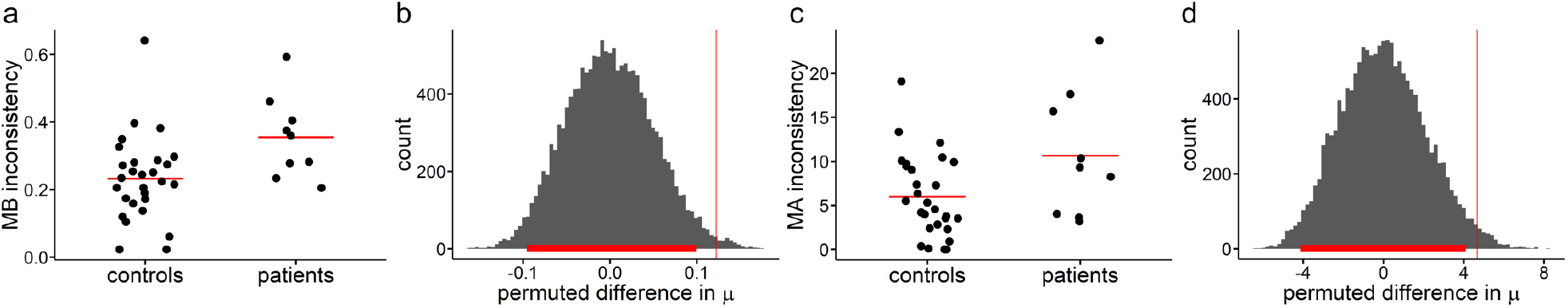
Within-participant changes in inter-temporal preferences. **a** and **c**, model-based T1-T2 choice inconsistency (deviation between T2 choices and model-predictions based on a model fitted to T1 data) and model-agnostic (MA) T1-T2 choice inconsistency (mean absolute deviation of indifference points between T1 and T2). **b** and **d** permutation test for model-based and model-agnostic inconsistency scores respectively. Histograms show null distributions of mean group differences across 10k randomly shuffled group labels; red vertical lines: observed group differences; red horizontal line: 95% highest density interval.

In the light of the slightly poorer model fit in patients vs. controls (see Table 3), which was reflected in overall lower softmax β parameters in patients (Figure 8A), three different control subgroups were selected based on different cut-off values of β (Figure 8A). Model-based choice inconsistency for a cut-off of *β* = .5 is shown in Figure 8B. Across three cut-off values, permutation tests again confirmed lower choice consistency in patients vs. controls (Figure 8C-E), illustrating that these effects are not driven by group differences in *β*.

**Figure 8.**
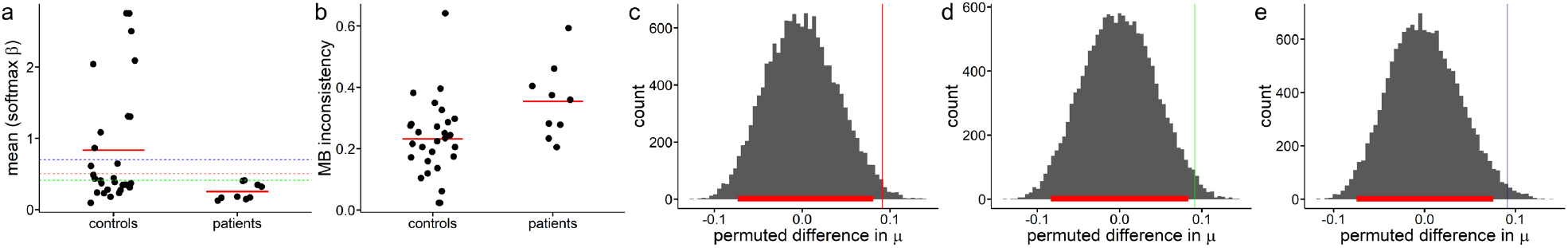
**a**, Individual-subject softmax *β* parameters (mean of T1 and T2 posterior medians) modelling decision noise. To account for effects of group differences in *β*, in additional analyses, controls matched on *β* to the patients were selected across various *β* cut-offs (*β=*0.5: red dotted line, n=17; *β*=0.41: green dotted line, n=14; *β*=0.7: blue dotted line, n=19). **b**, Individual-subject model-based T1-T2 choice inconsistency in *β*-matched controls (*β* cutoff = 0.5) and patients. **c-e**, Permutation test results for group differences in model-based choice inconsistency (histograms of group differences with 10k randomly shuffled group labels) across all cut-offs (**c:** *β=*0.5, **d:** *β=*0.41, **e:** *β=*0.7). In line with the analysis across the whole sample, patients showed significantly increased model-based inconsistency for all cut-offs (**c:** *p* = 0.018, **d:** *p* = 0.023, **e:***p* **=** 0.01). Vertical lines in c-e denote the observed group difference, horizontal red lines denote the 95% HDI of the null distribution.

## Discussion

We capitalized on the rare opportunity to longitudinally follow a group of patients with OCD receiving deep brain stimulation (DBS) to the anterior limb of the internal capsule / nucleus accumbens region (ALIC/NAcc), to examine chronic stimulation effects on temporal discounting. Crucially, according to current models of inter-temporal choice (Peters & Büchel, 2011) and self-control (Figner et al., 2010; Hare et al., 2009), the ventral striatum / nucleus accumbens regions plays a central role in this process in both rodents and humans (Bartra et al., 2013; Da Costa Araújo et al., 2009; McClure & Bickel, 2014; Pisansky et al., 2019), but direct evidence for a mechanistic role in human inter-temporal choice is lacking. Our data show that chronic DBS to the ALIC/NAcc region disrupts the stability of inter-temporal preferences, both reflected in group-level changes in reliability, and individual-level changes in preferences.

Control analyses fit in with previously reported increases in temporal discounting and elevated impulsivity in patients with OCD (Ong et al., 2019; Sohn et al., 2014), while contrasting in part with other studies that do not find any group differences between patients and controls (Carlisi et al., 2017; Norman et al., 2017; Steinglass et al., 2017). Thus, overall studies to date suggest that associations of OCD and intertemporal choice are heterogenous and possibly related to contributions of specific OCD subtypes or comorbidities (Pinto et al., 2014). Generative hierarchical Bayesian models then allowed us to estimate posterior distributions of test-retest correlations (Haines et al., 2020), separately for short-term and long-term stability, and for patients and controls. This revealed a reduction in long-term reliability of log[*k*] in patients undergoing DBS. Notably, this effect was robust to a number of control analyses, controlling for e.g. effects of the observed range of log[*k*] values, choice stochasticity effects, and the presence of OCD symptoms. Analyses of within-subject preference stability confirmed these results. Both model-based and model-agnostic measures confirmed a reduction in long-term preference stability in patients undergoing DBS compared to controls, and these analyses were again confirmed when groups were matched on choice stochasticity.

These findings suggests that, in addition to short-term plasticity processes (Kuhn & Baldermann, 2020), long-term ALIC/NAcc DBS (Denys et al., 2010) can interfere with the expression of intertemporal preferences that are thought to rely on this same circuitry (Hariri et al., 2006; Kable & Glimcher, 2007; McClure & Bickel, 2014; Peters & Büchel, 2011). While earlier reports noted effects of acute stimulation on risk-taking and impulsivity (Luigjes et al., 2011; Nachev et al., 2015) (albeit with acute block-wise stimulation protocols that differ substantially from our approach, where ON and OFF sessions were separated by 24h wash-out periods), our longitudinal analysis revealed changes only following prolonged stimulation and does not suggest a specific direction of change. Rather, the data might suggest a role of the ALIC/ NAcc region in maintaining preference stability over time. However, our data do not rule out the possibility that these effects are unspecific, in the sense that any chronic DBS protocol might have elicited the same effects. For example long-term DBS of the subthalamic nucleus is associated with increased plasticity, i.e. wide spread structural changes from sensory motor- to prefrontal cortex and limbic areas (van Hartevelt et al., 2014).

The exact cellular mechanisms underlying DBS effects remain speculative and poorly understood (Figee et al., 2013; Lozano & Lipsman, 2013; Robbins et al., 2019). Potential effects range from DBS acting as an informational lesion, to changes in inter-regional functional connectivity (Figee et al., 2013) and a general modulation of oscillatory activity and in consequence pathological circuitry (Lozano & Lipsman, 2013).

How might the NAcc region support preference stability in inter-temporal choice? First, it is well established that NAcc activity tracks subjective reward valuation across a range of different reinforcement learning and decision-making tasks and different classes of reinforcers (Bartra et al., 2013; Clithero & Rangel, 2014), including inter-temporal choice tasks (Basar et al., 2010; Kable & Glimcher, 2007; Peters & Büchel, 2011). Long-term DBS to this region might induce local plasticity, and thus disrupt or alter value representations in this circuit. Additionally (or alternatively), DBS might disrupt interactions of this circuit with other regions supporting inter-temporal choice, such as lateral prefrontal cortexand/or ventrialmedial prefrontal cortex. In line with this view, DBS of theNAcc has been shown to affect activation dynamics in frontostriatal circuits (Figee et al., 2013; Figee et al., 2014) and similar network interactions have been suggested to support successful self-control (Hare et al., 2009; Peters & D’Esposito, 2016).

A number of limitations of the present study deserve discussion. First, as noted above, the lack of a control group that received DBS to another target site limits the interpretability of our findings. Effects might reflect general DBS effects, rather than effects specific to the ALIC/NAcc region. However, given that DBS stimulation sites differ for different disorders, such studies would then require additional controls for differences in symptoms. Additional longitudinal studies are clearly required to clarify this. Second, we focused on inter-temporal choice, a stable decision trait with relevance for many mental disorders (Amlung et al., 2019; Lempert et al., 2019). Future work is required to establish whether the observed effects extend to other tasks or preferences. Third, we did not obtain neural read-outs such as functional magnetic resonance imaging data that might have allowed us to examine systems-level effects of DBS in individual patients (Figee et al., 2013; Smolders et al., 2013). The degree to which the observed effects are due to changes in neural systems beyond the stimulated target site therefore remain elusive.

In summary, the present findings yield an example for a subtle alteration in higher cognitive function following long-term (chronic) DBS to ALIC/NAcc. These data further reveal a potential contribution of this area to the maintenance of preference stability over time in the context of intertemporal choice that further studies might elaborate on.

## Acknowledgements

C.S. was funded under the Walter and Marga Boll Foundation (210-06-16) and the Institutional Strategy of the University of Cologne within the German Excellence Initiative (ZUK 81/1). JCB receives funding by the DFG (Project-ID 431549029 – SFB 1451). J.P. was supported by Deutsche Forschungsgemeinschaft (PE 1627/5-1).

## Author contributions

J.K., J.P. C.S. and B.W. designed the study. C.S., B.W. and M.M. acquired the data. C.S. and B.W. analyzed data. B.W. performed the modeling and statistical analyses. J.P. and J.K. supervised the project. B.W. and J.P. wrote the paper, and all authors provided revisions.

